# How Mg^2+^ stimulates DNA repair in prokaryotic (6-4) photolyases

**DOI:** 10.1101/366252

**Authors:** Hongju Ma, Daniel Holub, Natacha Gillet, Gero Kaeser, Katharina Thoulass, Marcus Elstner, Norbert Krauß, Tilman Lamparter

## Abstract

Prokaryotic (6-4) photolyases branch at the base of the evolution of cryptochromes and photolyases. In the *Agrobacterium* (6-4) photolyase PhrB, the repair of DNA with UV-induced (6-4) pyrimidin dimers is stimulated by Mg^2+^. We show that Mg^2+^ is required for efficient lesion binding and for charge stabilization after electron transfer from the FADH^-^ chromophore to the DNA lesion. Two highly conserved Asp residues close to the DNA binding site are essential for the Mg^2+^ effect. Simulations showed that two Mg^2+^ bind to the region around these residues. DNA repair by eukaryotic (6-4) photolyases is not increased by Mg^2+^. Here, the structurally overlapping region contains no Asp but positively charged Lys or Arg. During evolution, charge stabilization and DNA binding by Mg^2+^ was therefore replaced by a positive amino acid. We argue that this transition has evolved in a freshwater environment. Prokaryotic (6-4) photolyases usually contain an FeS cluster. DNA repair of a cyanobacterial member of this group which is missing the FeS cluster was also found to be stimulated by Mg^2+^.

## Introduction

Ultraviolet light induces the formation of additional bonds in two neighboring pyrimidine groups of DNA. Two major photoproducts, termed cyclobutane-pyrimidine dimer CPD- and (6-4) photoproduct, are distinguished. Without DNA repair, these additional bonds would lead to mutagenesis and cell death. The so-called nucleotide excision repair is present in almost all organisms. This kind of repair is based on the coaction of multiple enzymes and requires an intact complementary DNA strand as template [1]. The so called photorepair is based on the action of a a photolyase, which binds as a single protein to the DNA lesion. Photolyases are widely distributed in all domains of life, but missing on organisms such as higher mammals [2]. The two different DNA lesions, CPD and (6-4) photoproducts are repaired by two different subgroups of photolyases, termed CPD- and (6-4) photolyases. The central FAD chromophore of photolyases must be in the fully reduced state, FADH^-^, for repair activity. All photolyases share a light driven mechanism, photoreduction, during which oxidized FAD converts to FADH^-^. These electrons flow from the surface of the protein through a chain of three or four amino acid residues to FAD. After binding of the DNA lesion to a pocket of the protein, the small distance to FADH^-^ allows electron exchange. Excitation of FADH^-^ by light or by energy transfer from a photoexcited antenna chromophore triggers electron transfer to the lesion of the DNA. In a series of events, the nucleotides revert to the original state and the electron returns back to FADH.

Phylogenetically, 6 different groups of photolyases can be distinguished: class I CPD photolyases which comprises the founding member *Escherichia coli* photolyase together with mainly bacterial homologs, class II CPD photolyases with mainly eukaryotic members, the class III CPD photolyases, another group with mainly bacterial members, the CryDASH proteins, which are CPD photolyases that repair single stranded DNA, the eukaryotic (6-4) photolyases and a group of prokaryotic (6-4) photolyases that is termed FeS-BCP proteins [3]. The entire photolyase family contains also plant- and animal cryptochromes, flavoproteins that have lost DNA repair capabilities and serve as photoreceptors. These are phylogenetic sister groups of class III CPD photolyases and eukaryotic (6-4) photolyases, respectively [4].

Cryptochromes and photolyases share a common fold with primases [5], enzymes that synthesize RNA oligonucleotides during replication. According to structural alignments, the phylogenetic position of FeS-BCP proteins is between primases and other photolyases and cryptochromes. We therefore propose that the FeS-BCP proteins represent the most ancient group of photolyases and that comparative studies including FeS-BCP proteins lead to a better understanding of the early photolyase evolution [6]. Differences between FeS-BCP proteins and other photolyases that indicate evolutionary transitions are as follows:

(i) The two FeS-BCP proteins investigated so far, *Rhodobacter* CryB and *Agrobacterium* PhrB, have a DMRL antenna chromophore [6, 7]. Other photolyases and cryptochromes have either MTHF, 8-HDF or other antenna chromophores, but never DMRL. (ii) According to biochemical studies and sequence alignments, about 70% of FeS-BCP proteins including PhrB and CryB contain an iron sulfur (FeS) cluster. All other photolyase groups have no FeS cluster. Apparently, this cofactor was lost during evolution [3]. (iii) FeS-BCP members have a C-terminal extension, a feature that they share with the related cryptochromes and some other photolyases, although the sequences reveal no homologies between the different groups. The C-terminal extension of cryptochromes is relevant for signal transduction, and both CryB and PhrB could serve as photoreceptors for bacterial light responses [8, 9]. (iv) A loop between helix 7 and helix 8 (a7 and a8) of PhrB interacts with the DNA. This DNA interaction is replaced by another loop in other photolyases, e.g. the loop connecting helices a17and a18 in *Drosophila* (6-4) photolyase [10]. (v) The photoreduction chain is composed of two Trp and one central Tyr in FeS-BCP proteins. This function was overtaken by an unrelated Trp triad in other photolyases and cryptochromes [11] (vi) The repair activity for (6-4) photoproducts of short single stranded oligonucleotides by PhrB and CryB was increased about thousandfold by the addition of Mg^2+^ or other divalent cations [12]. Such an effect is absent from photolyases of other groups. The early diversification of photolyases is thus characterized by multiple significant changes that indicate severe evolutionary pressure and general adaptation processes.

In the present work we investigate the induction of DNA repair by Mg^2+^. A proposed Mg^2+^ binding position next to the DNA lesion was modified by site directed mutagenesis. The inductive effect of Mg^2+^ was lost for both mutants investigated, although the base repair activity without Mg^2+^ remained unaffected by the mutations. We also investigated the Mg^2+^ effect for different lengths of single stranded and double stranded DNA. The action of Mg^2+^ was also analyzed by computational MD approaches. These studies revealed a high probability of two Mg^2+^ next to the proposed positions. Our results show that Mg^2+^ increases the binding affinity for DNA and stabilizes the negative charge on the DNA after electron transfer from FADH^-^ to the lesion. We discuss by which events the loss of Mg^2+^ dependency could have been driven during evolution and how the positive charges of Mg^2+^ were replaced.

## Results

### Repair of (6-4) lesions by PhrB, different lengths of single stranded oligonucleotides, double stranded oligonucleotides and Mg^2+^ effect

The repair activities of PhrB and CryB, which belong to the group of prokaryotic (6-4) photolyases or FeS-BCP proteins, are drastically increased by the addition of Mg^2+^, whereas the repair activities of photolyases belonging to all other groups are not affected by divalent cations [12]. In order to study the effect of Mg^2+^ more in detail, we intended to compare repair rates of different lengths of oligonucleotides. In previous work [6, 12], the oligonucleotide "t_repair" with 8 nucleotides was used. Here we initially compared the PhrB repair activities for single stranded oligonucleotides with 8, 10, 12 and 15 bases in the absence of Mg^2+^ (Fig. 1). No repair of the 8-mer "t_repair" was detectable by HPLC for irradiation times of up to 40 min and 60 min irradiation resulted in a repair of only (0.9 ±0.1) %. This very low percentage in comparison to earlier work is due to lower protein / DNA ratio. For the (6-4) photoproduct of the 10-mer ODN4, there was no detectable signal of repaired DNA for illumination times up to 20 min, and after 40 min irradiation (4.2±0.2) % were repaired. With (6-4) photoproducts of ODN5 and ODN6, which have 12 and 15 oligonucleotides, respectively, about half of the lesion DNA was repaired after 20 min and the repair was complete after 40 min of irradiation. We concluded that the weak repair activity observed in our previous studies [6] was not only due to the lack of Mg^2+^ but also due to the short length of the oligonucleotide. The more efficient repair with longer oligomers results most likely from an increased binding of the oligomers to PhrB. Because there was no increase between 12-mer and 15-mer, we assume that there would be no further improvement for larger oligomers.

**Fig. 1.**
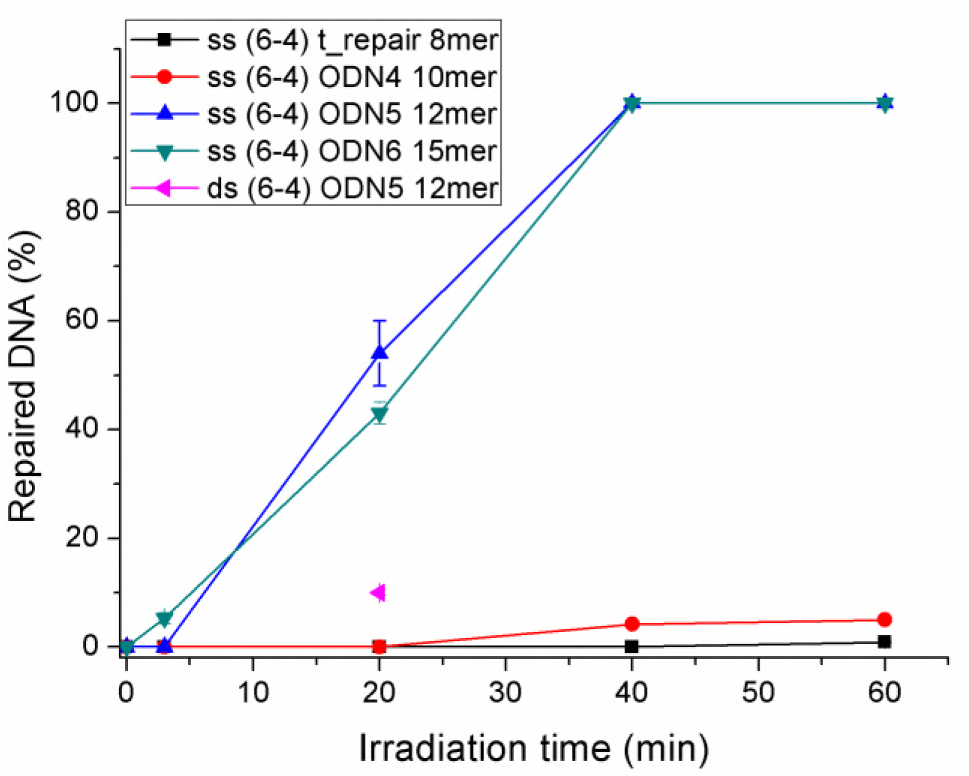
Repair of single stranded (6-4) photoproducts of different lengths and ds (6-4) ODN5 by the photolyase PhrB. The repair assays were performed in the absence of divalent cations. Proportions of repaired DNA of (6-4) photoproducts of single stranded DNA oligonucleotides "t-repair" (8-mer, black squares), ODN4 (10-mer, red circles), ODN5 (12-mer, blue up-pointing triangles), ODN6 (15- mer, down-pointing triangles) and double stranded ODN5 (12-mer, pink left-pointing triangles) are shown as a function of irradiation time. Irradiation times were 0, 3, 20, 40 and 60 min. Mean values ± SE from 3 independent experiments. In those cases where error bars are invisible, the errors are smaller than the symbols. Example HPLC profiles are shown in Fig. S1.

The same repair assay was performed with double stranded DNA. To this end, the ODN5 photoproduct was mixed with its complementary oligonucleotide, heated to 90 °C and cooled down over a period of 3 h. The melting temperature (T_m_) of double stranded ODN5 is 10 °C higher than the reaction temperature of 22 °C such that the DNA would form stable double strands. After 20 min incubation with PhrB, the yield of repaired DNA was only ca. 20% of that found for the single stranded ODN5 after the same period of time (Fig. 1 and Table 1). Thus, single stranded (6-4) DNA is more efficiently repaired than double stranded DNA.

**Table 1.** Repair of single and double stranded oligonucleotides of different lengths by PhrB. DNA and protein concentrations were 5 μM and 0.5 μM, respectively. REA: relative enzyme activity; concentration of repaired DNA divided by protein concentration and time. Mean values of 3 or more independent experiments ± SE. The "fold increase" is the RAE with Mg^2+^ divided by the RAE without Mg^2+^

| Oligo DNA | Length (nt) | ss or ds substrate | $Mg^{2+}$ | Irradiation time | Repair (%) | REA ($min^{-1}$) | fold increase |
| --- | --- | --- | --- | --- | --- | --- | --- |
| t-repair | 8 | ss | - | 2 h | $3.5 \pm 0.3$ | $0.3 \pm 0.03$ | |
| t-repair | 8 | ss | 4 mM | 3 min | $54 \pm 1$ | $180 \pm 4$ | 620 |
| ODN5 | 12 | ss | - | 20 min | $54 \pm 6$ | $27 \pm 3$ | |
| ODN5 | 12 | ss | 4 mM | 2 min | $81 \pm 2$ | $405 \pm 10$ | 15 |
| ODN5 | 12 | ds | - | 20 min | $10 \pm 0.5$ | $5 \pm 0.3$ | |
| ODN5 | 12 | ds | 4 mM | 2 min | $35 \pm 0.5$ | $175 \pm 3$ | 35 |

The repair assays were performed with PhrB in which the FAD chromophore was initially in the oxidized state. It is generally assumed that DNA repair is only possible with reduced FADH^-^. In our assays the light treatment during co-incubation triggers two light-dependent processes, photoreduction of FAD [13] and electron transfer from FADH^-^ to the DNA lesion. The fact that 2 min co-incubation in the light resulted in significant repair of (6-4) DNA (Fig. 1 and later experiments) shows that photoreduction during this short irradiation time is sufficient to generate active reduced FADH^-^. In control experiments we tested whether 60 min pre-irradiation would increase the repair activity. When the protein/DNA incubation time was 2 min, pre-irradiation resulted in a ca. 30% higher repair. When the protein/DNA incubation time was 20 min or longer, preirradiation had no significant effect.

In subsequent comparative experiments we most often chose conditions in which the yields of repaired DNA were lower than 100% but large enough to recognize the peak of repaired DNA clearly in the HPLC analysis. Usually, this was achieved by variations of incubation time between 2 and 120 min. For semi-quantitative comparisons we calculated a relative enzymatic activity (REA) by dividing the concentration of repaired DNA by the concentration of the protein and the incubation time. According to the above control measurements, activities measured with short incubation times are underestimated by up to 30%, which is, however, irrelevant for the subsequent comparisons. To assume a linear relationship between the yield of repaired DNA and incubation time is justified based on the results with ODN5 shown in Fig. 1, whereas a monoexponential decay, as proposed in earlier studies [12], is not observed: the time dependence of lesion repair (Fig. 1) could not be fitted with a monoexponential function.

We then tested the increase of repair by Mg^2+^ for (6-4) photoproducts of single stranded 8-mer t_repair, single stranded 12-mer ODN5 and double stranded 12-mer ODN5. As given in Table 1, the REA increased 620 times for the 8-mer t_repair, 15 times for single stranded 12-mer ODN5 and 35 times for double stranded ODN5. Thus, for all constructs, the repair was improved dramatically by Mg^2+^ but the relative effect was lower if the repair without Mg^2+^ was already high, as *e.g.* in the case of single stranded ODN5. We propose that the large increase for short fragments is partially due to a role of Mg^2+^ in DNA binding. Longer DNA is expected to bind stronger even without Mg^2+^. For longer DNA we therefore propose that Mg^2+^ has an independent effect such as charge stabilization.

### DNA repair by the PhrB-Y424F mutant

The PhrB-Y424F mutant was previously generated to test for possible electron transfer between the iron sulfur cluster of PhrB and the DNA lesion [14]. According to a DNA binding model, Tyr424 interacts directly with the DNA lesion [13]. In the mutant this binding was found to be very weak, the k_D_ value for a 15-mer oligonucleotide with (6-4) lesion was 18 μM, whereas for wild type PhrB, a k_D_ of 13 nM was estimated [14]. In the previous study with PhrB-Y424F, no repair activity was found [14]. The crystal structure of PhrB-Y424F showed that the folding of the protein is unaffected by the mutation and effects can be directly assigned to the replacement of this amino acid [13]. In order to test for a Mg^2+^ effect in this mutant, we performed repair assays with protein concentrations of 5 μM, *i.e.* 10 times higher than in the other experiments presented here. We concentrated on the single stranded ODN5 substrate, which is most efficiently repaired by PhrB, and found indeed that even in the absence of Mg^2+^ a small fraction of DNA was repaired by PhrB-Y424F (Fig. 2 for HPCL). The REA value of (5±1) x 10^−4^ min^-1^ was drastically lower than for the wild type, as expected. In the presence of Mg^2+^, the REA value was 0.2, thus 420 times higher than for the repair without Mg^2+^ (Table S2). This increase is thus significantly larger than the increase for wild-type PhrB with the same oligonucleotide. We assume that the stronger effect as observed for the mutant is based on a larger improvement of DNA binding by Mg^2+^.

**Fig. 2.**
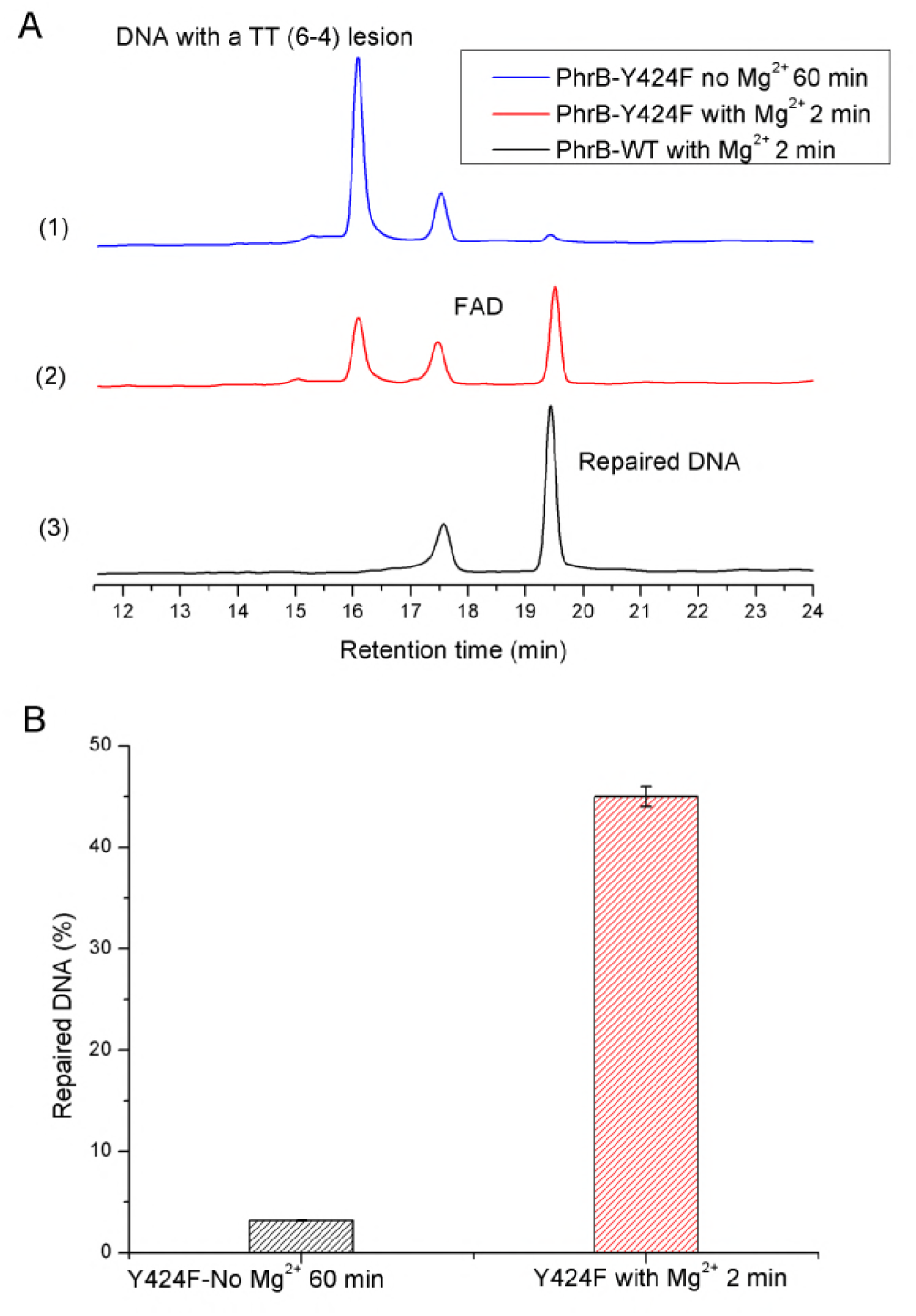
DNA repair activity of PhrB wild-type and the mutant PhrB-Y424F with and without Mg^2+^. Repair of single stranded ODN5 photoproduct. DNA concentration in all assays 5 μM, protein concentration in Y424F assays 5 μM and in PhrB wild type assays 0.5 μM (A) examples for HPLC profiles. (B) Proportion of DNA repaired by PhrB-Y424F in the absence and presence of Mg^2+^.

### DNA repair by the PhrB-D179N and PhrB-D254N mutants

Based on our results we expected that the active Mg^2+^ ions in PhrB would be close to the DNA lesion. Because the Mg^2+^ effect is only found in PhrB and the related CryB, but not in other groups of photolyases [12], the most probable site for Mg^2+^ action seemed to be at a specific α7-α8 connecting interdomain loop which is only present in PhrB and its relatives, and which is replaced by the α17-α18 connecting loop in other photolyases [6]. In PhrB, the loop contains 4 negatively charged amino acids, Asp179, Glu181, Asp189, and Asp201. Of these, only Agp179 is highly conserved in the FeS-BCP proteins. Another nearby Asp at position 254 has its carboxylate side group ca. 7 Å distant from Asp179 and is also highly conserved in FeS-BCP proteins [12]. We found that even in a set of 4000 BLAST homologs, both amino acids are conserved. Both residues seemed to be the most likely candidates for Mg^2+^ binding close to the DNA lesion. To test this hypothesis, we generated both relevant mutants, PhrB-D179N and PhrB-D254N in which the negatively charged side chain of each Asp was replaced by a neutral side chain of Asn. Both mutants were used for repair assays with and without Mg^2+^ (Fig. 3). In these repair assays, the (6-4) photoproduct of t_repair was used as the substrate. Without Mg^2+^ and during a 2 h irradiation, wild type PhrB, PhrB-D179N and PhrB-D254N repaired (3.5 ± 0.3) %, (4 + 0.1) %, (5 + 2) % of the DNA, respectively. When Mg^2+^ was added and the reaction mixture irradiated for 3 min, PhrB repaired (54+ 1) %, but for PhrB-D179N and PhrB-D254N no signal from repaired DNA was detectable. The irradiation time was therefore prolonged to 2 h. For each mutant, the yields of repaired DNA were almost the same no matter if the repair assays were conducted with or without Mg^2+^ (Fig. 3). Hence, Asp179 and Asp254 are crucial for enhancement of the catalytic activity of PhrB by Mg^2+^, which is likely due to binding of the divalent cation to these residues during the reaction cycle of DNA repair.

**Fig. 3.**
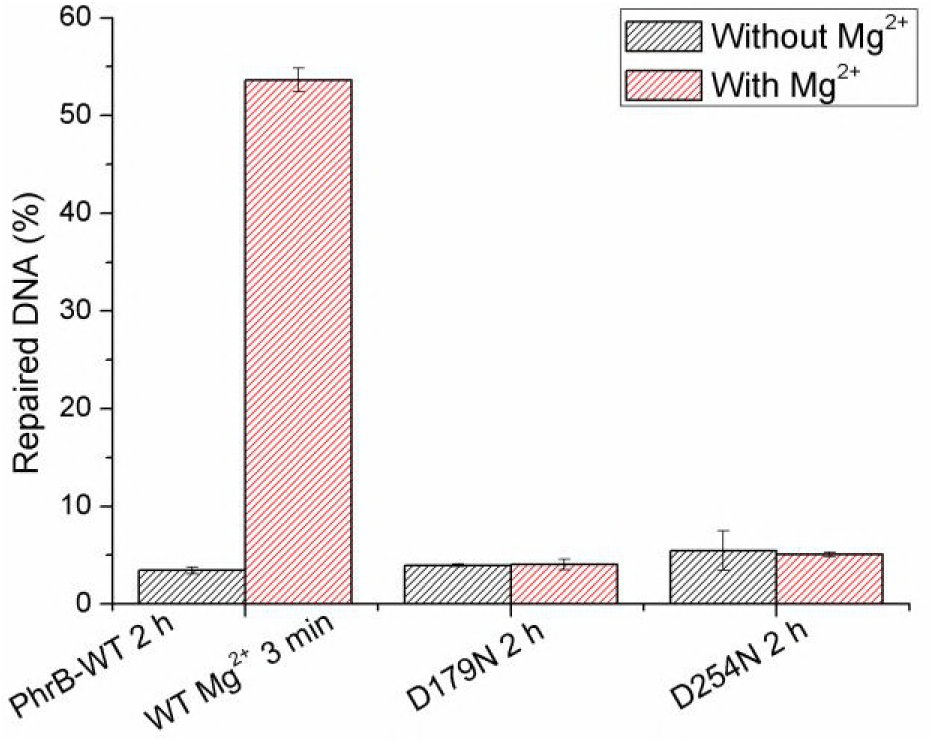
DNA repair of PhrB-WT and its mutants PhrB-D254N and PhrB-D179N with and without Mg^2+^. 8**-**mer single stranded t_repair photoproduct was used in the repair assays. DNA and protein concentrations were 5 μM and 0.5 μM, respectively. Irradiation times for PhrB-WT without and with Mg^2+^ were 2 h and 3 min, respectively; for the two mutants PhrB-D179N and PhrB-D254N it was always 2h. Mean values ± SE of 3 independent measurements.

#### A cyanobacterial photolyase without FeS cluster

About 30% of the proteins that belong to the group of FeS-BCP proteins lack the four FeS binding Cys residues and have therefore no FeS cluster. As a representative of this subgroup we expressed a putative (6-4) photolyase (accession number WP_011132061) of the marine cyanobacterium *Prochlorococcus marinus.* This protein is termed PromaPL here. According to phylogenetic studies, PromaPL belongs to the FeS-BCP clade. The expression levels in our *E. coli* based recombinant system were very low, despite several rounds of optimization. After affinity chromatography the protein could be purified to only ca. 10% of the co-purified protein (Fig. S2). Since *E. coli* has no (6-4) photolyase, any (6-4) repair activity of this preparation can however be assigned to the partially purified PromaPL. Using single stranded ODN5 as substrate, we observed a clear repair activity in the presence of Mg^2+^, whereas without Mg^2+^, no repair was found. Thus, the Mg^2+^ effect is also present in prokaryotic (6-4) photolyases without FeS cluster (Fig. 4). The RAE of PromaPL in the presence of Mg^2+^ (0.012 min^-1^) was much smaller than that of PhrB (405 min^-1^, see legend of Fig. 4). The FAD chromophore is recognized in the HPLC profiles of the repair assays, indicating a ca 10x lower FAD incorporation in PromaPL than in PhrB on a molar basis. However, this does not account alone for the huge RAE difference. We consider that the low activity could directly or indirectly (via inappropriate protein folding) be due to the loss of the FeS cluster in PromaPL.

**Fig. 4.**
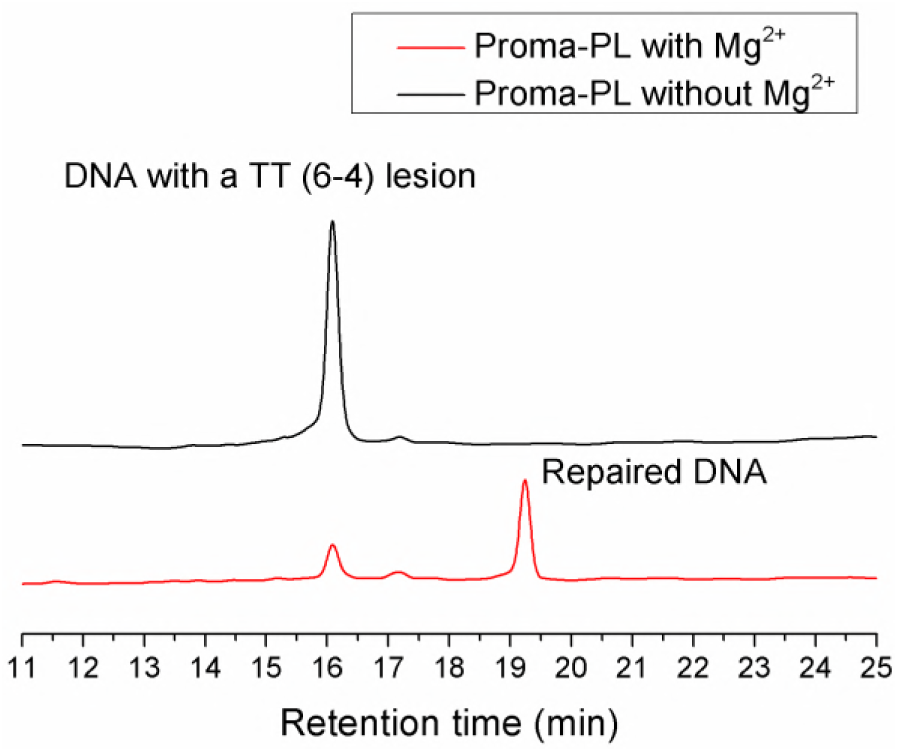
Repair of single stranded ODN5 photoproduct by *Prochlorococcus marinus* photolyase PromaPL. incubation time 60 min. Examples of HPLC profiles. Upper panel: repair reaction without Mg^2+^, lower panel: repair reaction with 1.5 mM Mg^2+^. The irradiation time was 60 min. DNA and PromaPL concentrations were 5 μM and 5 μM, respectively. In the absence of Mg^2+^, no repair was detectable (3 assays). In the presence of 1.5 mM Mg^2+^ the RAE was 0.012±0.0009 min^-1^ (mean of 3 experiments ± SE)

#### Molecular dynamics studies on Mg^2+^ binding

In order to get deeper insight into the binding sites and the number of bound Mg^2+^ close to the active site of PhrB, we performed molecular dynamics simulations including the whole solvated protein, a double stranded damaged 12 nucleotides long DNA and Mg^2+^ cations. The DNA and its interaction mode with the protein were taken from the *Drosophila* (6-4) photolyase cocrystal structure [10] and fitted to the PhrB structure by slight rotations of nucleotide residues to avoid steric clashes. Molecular dynamic simulations were always performed for 1 μs after adding Mg^2+^ to the system.

In the active sites of the *Drosophila* (6-4) photolyase cocrystal and of our PhrB/DNA structure, the (6-4) photoproduct interacts with His366 (PhrB numbering). The PhrB-H366N mutant shows no repair activity [14]. Several studies on (6-4) photolyases suggest that a proton transfer occurs from this histidine residue to the damaged DNA as a part of the repair process [15-18]. The His366 is expected to be protonated in the ground state and transiently deprotonated for 10 ns or longer during the repair process [15], but also a deprotonated His366 in the ground state and another reaction mechanism could be possible in PhrB. For our simulations on the localization of Mg^2+^ we consequently carried out two sets of simulations, one with positively charged His366^+^ and one with a neutral His366.

In PhrB-WT with protonated His366^+^, we observe after few nanoseconds upon onset of our simulations the stable binding of two Mg^2+^ ions close to the (6-4) photoproduct and close to Asp179 and Asp254 (see Fig. 5 and 6). One ion, termed 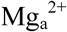, is close to the phosphate group of the 3’ thymine and interacts with Asp179 and Glu181. The second ion, termed 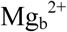, is deeper buried in the active site and interacts with the phosphate of 3’ thymine, the oxygen atom of the 3’ thymine ring and with Asp254 (Table 2).

**Fig. 5.**
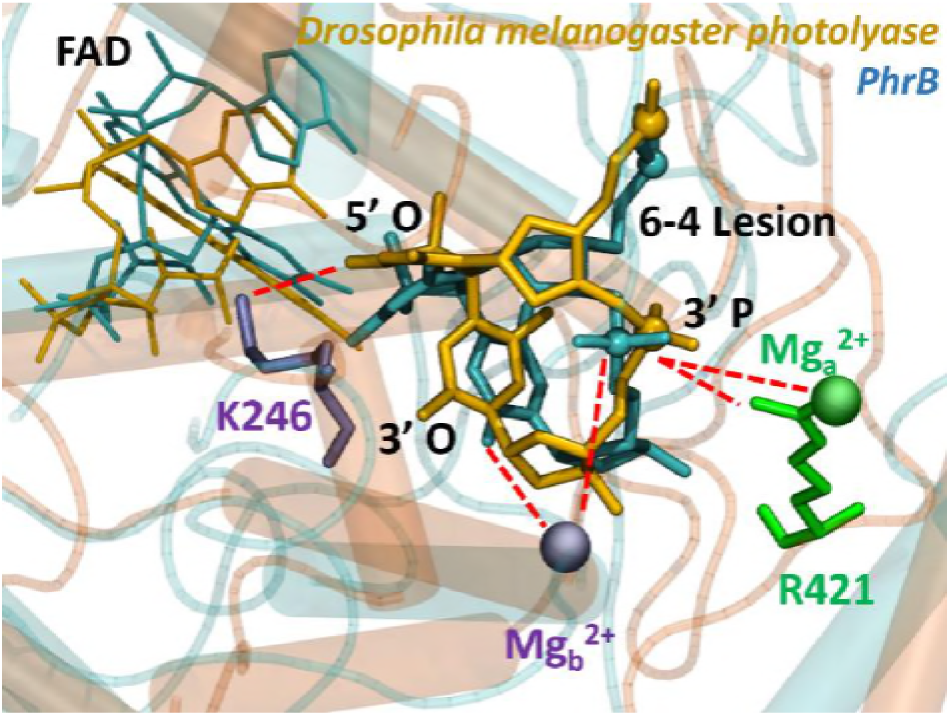
Comparison between PhrB (cyan, MD snapshot) and *Drosophila melanogaster* photolyase (orange PDB 3CVU) [10] active site containing the 6-4 photoproduct (shown as sticks, phosphor atoms are represented by balls), FAD (smaller sticks), and positively charged groups: R421 (from *Drosophila melanogaster* photolyase) and equivalent 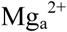 (from PhrB) in green; K246 (from *Drosophila melanogaster* photolyase) and equivalent 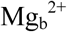 (from PhrB) in violet. The possible electrostatic or hydrogen bonding interactions are represented by dashed red lines. It has been shown previously by MD simulations that K246 can interact with the 3’ phosphate group P atom [19].

**Fig. 6.** Positions occupied each nanosecond by magnesium cations along the 1 μs simulations in PhrB with positively charged or neutral His-366, D179N or D254N mutants. The cations in position 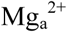 and 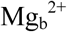 are represented by green and purple spheres respectively. Amino acid residues and protein backbone are taken from a single snapshot along the simulation.

**Table 2.**
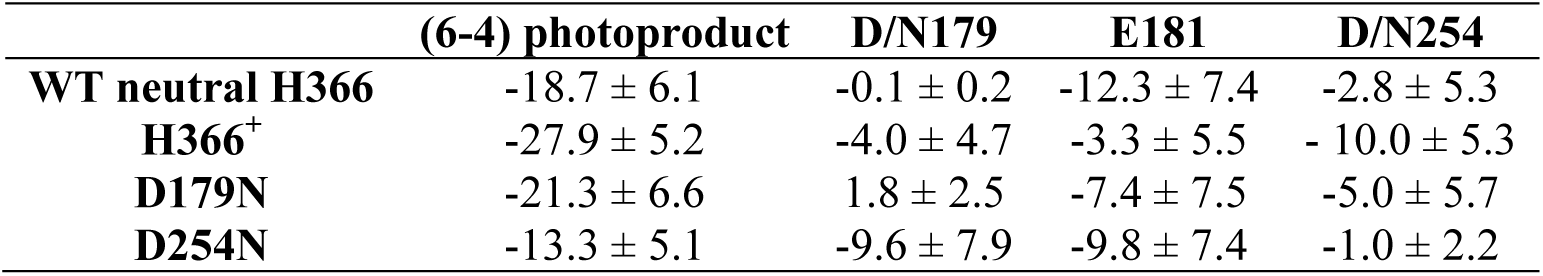
**Electrostatic interactions** in kcal/mol between Mg^2+^ cations and (6-4) photoproducts or acid residues during molecular dynamic simulations. Averaged electrostatic interactions between the Mg^2+^ cations and the (6-4) photoproduct or D179, E181, D254 or the corresponding asparagine after mutation. Negative sign means attractive electrostatic interactions while positive sign refers to electrostatic repulsion. The error values correspond to the standard deviation of the interactions during the 1 microsecond molecular dynamic simulations.

In both PhrB-D179N and PhrB-D254N mutants, the stability of the binding sites of the Mg^2+^ cations close to the (6-4) photoproduct is affected by the lack of the aspartate negative charges (Table 2). In the PhrB-D254N mutant, the position at 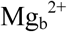 is no longer occupied while the 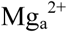 is present, interacting even stronger with Asp179 or Glu181 than in wild-type PhrB (Table 2). The PhrB-D179N mutant presents an unexpected behavior: the 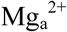 position is occupied all along the 1 μs MD simulation, partially stabilized by an interaction with Glu181, while 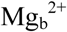, present at the beginning, is released after 500 ns. The simulation results are in agreement with the experiments with the D254N and D179N mutants in which the Mg^2+^ effect on DNA repair was lost.

With neutral His366 in PhrB we observe a slightly different conformation of this side chain as compared to protonated His366^+^, and the binding of only one Mg^2+^ in the active site, close to the 3’ thymine (Fig. 6). The cation, however, mainly interacts with Glu181 (Table 2), whereas the electrostatic interactions with Asp179 or Asp254 are very small and underline the absence of strong ionic interaction between Mg^2+^ and these two aspartates. Actually, Asp254 interacts with Arg187 during the first 750 ns of the simulation while Asp179 presents strong hydrogen bonds with Arg183.

A structural comparison with eukaryotic (6-4) photolyase from *Drosophila melanogaster* showed that these two aspartate positions are closely related to Arg421 and Lys246, respectively (Fig. 5). It has been reported that these side chains have a key role in binding of lesion DNA [10, 19]. At the position of Arg421, most of the homologs of *Drosophila* (6-4) photolyase have a histidine residue, which can be neutral or positively charged at physiological pH. The positive charge at the Lys246 spatial position is highly conserved in eukaryotic (6-4) photolyases. According to our PhrB simulations, 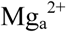 interacts with phosphate groups of the DNA in the periphery of the active site and should help to maintain the lesion in an optimized conformation to facilitate its flipped-out conformation. The presence of a positive charge in direct interaction with the damaged thymines would increase the electron affinity of the (6-4) lesion. In the crystal structure, the Lys246 side chain is orientated to form hydrogen bonds with the 5’ ring whereas 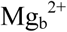 interacts with 3’ ring. Nevertheless, recent molecular dynamics simulations of *Drosophila melanogaster* photolyase have shown a motion of the lysine toward the 3’ phosphate group and thus closer to the 3’ thymine ring [20] which is in better agreement with our Mg^2+^ binding conformation. PhrB-D179N and PhrB-D254N mutations affect the 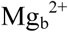 complex in our simulations, which is consistent with a weaker stabilization of the negatively charged (6-4) photoproduct intermediate during the repair catalytic cycle than in PhrB-WT.

## Discussion

When the PhrB repair of DNA carrying a (6-4) lesion was tested with oligonucleotides containing 8 to 15 bases, the best repair rate was obtained with the 12-mer and no further increase was obtained with the 15-mer. The repair of double stranded DNA on the other hand was less efficient as compared to the single stranded counterpart. Under improved reaction conditions, a weak repair activity was even obtained with the Y424F mutant, for which no repair had been observed in previous assays. The relative Mg^2+^ effect was strong (620 fold) for short oligonucleotides and weaker (22 fold) for longer oligonucleotides. With PhrB-Y424F and the long oligonucleotides, also a strong increase of repair activity by Mg^2+^ was observed. These length-dependent and binding-site dependent effects show that the increase of repair activity by Mg^2+^ is partially due to an improved binding of DNA to the protein. The PhrB-D179N and PhrB-D254N mutant results confirmed that Mg^2+^ acts close to the DNA binding site and the molecular dynamics simulations showed that under standard conditions two Mg^2+^ are bound to the relevant region of the protein. A direct interaction of protein bound Mg^2+^ with a phosphate of the lesion DNA is also evident from the model calculations. Besides improved binding, a second effect of Mg^2+^ became evident: even if optimum binding of lesion DNA is achieved, the DNA repair is still substantially increased by Mg^2+^. This second effect is the charge stabilization during repair. For both CPD- and (6-4)-photolyases the repair is initiated by electron flow from FADH^-^ to the lesion, in case of (6-4) photoproduct the electron is initially located on the 5' thymine at position 3, 4 or 5 [21, 22]. In the quantum chemical model with His366^+^ the distances between the Mg^2+^ ions and the isoalloxazine center of or the adenine center of FAD are 15 - 17 Å, whereas the distances between Mg^2+^ and the carbon 4 of the lesion thymine are shorter, i.e. 8-9 Å (Fig. 5 and distance measurements on the model). Therefore, the positive charges of both Mg^2+^ direct the electron of excited FADH^-^ towards the DNA lesion and increase the electron affinity of the lesion thymine. Without Mg^2+^ the lesion is less appealing for the excited electrons from FADH^-^ and inefficient back electron transfers become more likely. Although more studies are required to get detailed information of this topic, such a model can explain the low quantum yield of photorepair that is observed in the absence of Mg^2+^. The electron transfer is most likely followed by a rapid proton transfer from His366^+^ to the lesion DNA. According to our calculations, the change from positive to neutral His366 results in the loss of a Mg^2+^ in the surrounding of the lesion binding site. The distances to the lesion and to the FADH ring systems remain similar, but now two positive charges are lost. This will facilitate electron back transfer from the lesion to FADH. In summary, the Mg^2+^ system could have a beneficial effect on the DNA repair by PhrB in three ways, (i) coordination of the lesion within the protein pocket and DNA binding, (ii) charge stabilization after electron transfer from FADH^-^ to the lesion and (iii) support of electron back transfer from the lesion to FADH upon deprotonation of His-366.

In a previous study, the Mg^2+^ effect on DNA repair was compared for different photolyases. Such an effect was observed for FeS-BCP and another member of the FeS-BCP proteins, CryB [12]. In photolyases of other groups there was no such Mg^2+^ dependency. In addition, the standard reaction buffers in which photorepair is studied do not contain Mg^2+^ (e. g. [23]). Thus, two different metals were lost during the evolutionary transition from FeS-BCP proteins to other photolyases, iron and magnesium. In a new member of the FeS-BCP group, a photolyase from *Prochlorococcus marina*, which has no FeS cluster, we also found a stimulation of DNA repair by Mg^2+^. The loss of iron thus occurred more often in the evolution of photolyases than the loss of magnesium (Fig. 7). Iron often reaches limiting concentrations in the environment. In the oceans the iron concentrations range from 2 nM to 1 μM, depending also on biological activity. Iron fertilization in the oceans leads to algal blooms [24]. Iron loss could therefore have evolved in the oceans. Iron-free PhrB homologs are mainly found in cyanobacteria. The evolution of this branch of photolyases is thus consistent with such a scenario. However, the loss of the magnesium effect as observed for the major group of photolyases and cryptochromes cannot have taken place in the oceans, because marine Mg^2+^ concentrations are always in the range of 50 mM and there is no indication for significant losses due to biological activity [25]. Essentially comparable Mg^2+^ concentrations can be expected to have been present in the oceans even 2 Bio years back, before the evolution of eukaryotes began. At this time (or even earlier) the group of FeS-BCP and the other photolyases / cryptochromes must have diverged. We would therefore exclude the oceans as place of the separation of the major group of photolyases and cryptochromes from the FeS-BCP proteins. We also do not see an advantage of repair activity accompanied by the loss of Mg^2+^ stimulation. The comparison of PhrB (with Mg^2+^) with a eukaryotic photolyase has shown that the repair activities are even slightly higher for PhrB [12].

**Fig. 7.**
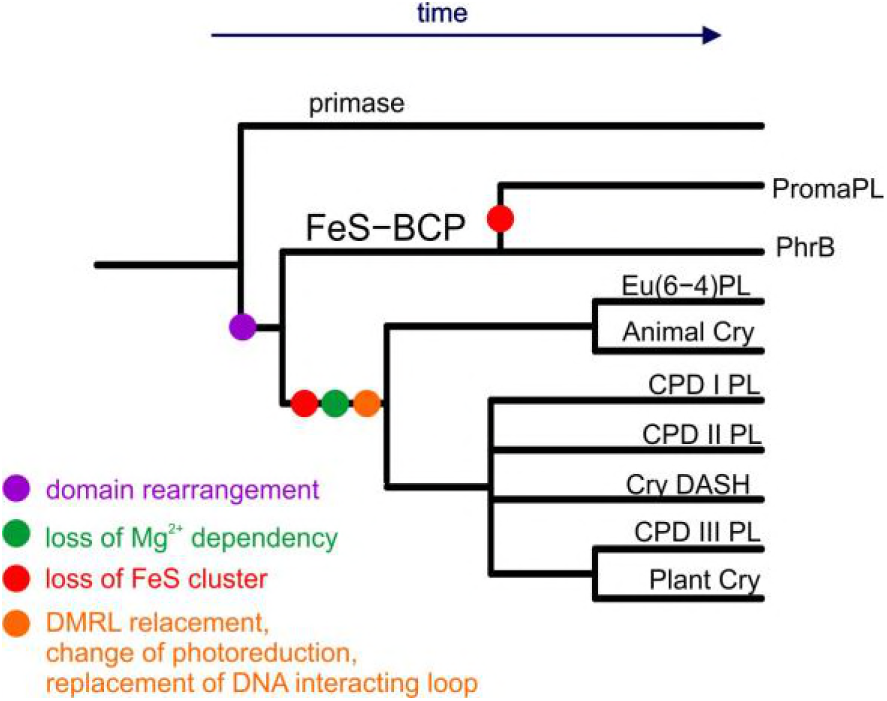
Evolution of photolyases and cryptochromes. A common fold in the large subunit of archaeal and eukaryotic primase, PriL, and photolyases / cryptochromes together with the FeS cluster in PriL and FeS-BCP proteins shows that the first branch point in the photolyases / cryptochrome family is between the FeS-BCP and the other groups. The loss of the FeS cluster occurred twice, within the group of FeS-BCP proteins and before evolution of the photolyases and cryptochromes that do not belong to FeS-BCP. This transition is also characterized by multiple other changes. The loss of Mg^2+^ dependency occurred probably only once.

However, freshwater lakes or rivers as well as rainwater pool biotopes could contain very low concentrations of magnesium. From our present view it seems that DMRL replacement, FeS-cluster-loss, Mg^2+^-activation-loss, replacement of the DNA interacting loop, replacement of the photoreduction chain occurred in a series of changes that were completed in the founding member of the major branch of photolyases and cryptochromes. The Mg^2+^ effects suggest that the founding member derived from a freshwater species or a descendant thereof.

## Materials and Methods

### Mutagenesis and expression

The PhrB mutants D179N and D254N were obtained by Quik Change site-directed mutagenesis (Agilent). Expression and purification of PhrB and the mutants were performed as described earlier [26]. The final buffer solution was 50 mM Tris/Cl, 5 mM EDTA, 300 mM NaCl, 10% glycerol, pH 7.8.

### Generation of (6-4) photoproducts

The used oligonucleotide sequences [27] are listed in Table S1. A defined concentration of the oligonucleotide, dissolved in water, was irradiated with UV-C. The (6-4) photoproduct was purified by HPLC [28]. Double stranded ODN5 was obtained by mixing (6-4) ODN5 and its complementary oligonucleotide, heating to 95 °C followed by slow cooling to 25 °C.

### Repair assay

The photorepair reaction mixture contained 5 μM of the (6-4) photoproducts in repair buffer (50 mM Tris-HCl, pH 7.5, 1 mM EDTA, 100 mM NaCl, 5 % (w/v) glycerol, 14 mM DTT). After 5 min pre incubation in darkness at 22 °C, aliquots were irradiated with 400 nm light emitting diodes (250 μmol m^-2^ s^-1^) for the indicated time. After separation of protein from DNA, the DNA was subjected to HPLC. The repair efficiency was estimated from the peak areas corresponding to damaged DNA and repaired DNA as described earlier [14].

### Photolyase from *Prochlorococcus marinus*

PhrB homologs [6] were screened for sequences without FeS-binding cysteines. We selected a sequence from *Prochlorococcus marinus* ssp CCMP1986 [29] (Uniprot identifier or the protein: Q7V2P7) and synthesized a codon optimized gene (Genscript) for expression in the pET28a vector (Novagen). The expressed gene has an N-terminal His-tag. Protein was expressed in *E. coli* ER2566 in autoinduction medium [30]. The final buffer was as for PhrB.

### Computational Methods

For the PhrB-DNA model the PhrB WT structure (PDB 4DJA) [6] and the damaged DNA bound to *Drosophila melanogaster* (6-4) photolyase (PDB 3CVU) [10]were used. Three starting structures have been defined considering the three possible protonation states of H366: Nεs protonation, Nδ protonation or positively charged and double protonated histidine. The force field parameters for DMRL and the iron sulfur cluster have been described earlier [11]. The FADH-parameters were taken from the AMBER force field [31, 32] and the charges were determined using the RESP approach [33]. All simulations were performed with the GROMACS 2016 package [34, 35] using the AMBER-SB99-ILDN force field [31, 32].

## ACKNOWLEDGEMENTS

This work was supported by the China Scholarship Council (H.M.). N. Gillet thanks the Alexander von Humboldt Foundation for fundings. Computational resources were provided by the state of Baden-Württemberg through bwHPC.

Author contribution: Wrote the paper: HM, NG, NK, TL; performed PhrB experiments: HM, KT, performed PromaPL experiments: HM, GK; performed MD calculations: DH, NG, ME; coordinated work: HM, NG, ME, NK, TL

The authors declare that they have no conflict of interest.

## Supplemental Information

### Supplemental Methods

#### Mutagenesis and expression

For expression of PhrB, an *E. coli* strain ER2566 carrying a pET21b based PhrB expression vector was used as template DNA [26]. The PhrB mutants D179N and D254N were obtained by site-directed mutagenesis according to the Quik Change site directed mutagenesis protocol (Agilent). The following primers were used; the triplet of the mutation is printed in bold:

D179N, GGCGGGCGCTGGAATTTT**AAT**GCGGAGAACCGCCAACCC and

GGGTTGGCGGTTCTCCGC**ATT**AAAATTCCAGCGCCCGCC;

D254N, GGCGCCACGCAG**AAT**GCCATGCTGCAGGATGAC and

GTCATCCTGCAGCATGGCATTCTGCGTGGCGCC.

For PCR amplification, the Q5 DNA polymerase (NEB) was used. Expression and purification of PhrB and the mutants were performed as described earlier for wild type PhrB [26]. In brief, protein with a N-terminal His-tag was expressed in *E. coli* strain ER2566 at 14 °C in LB medium; specific induction was induced by 0.1 mM IPTG. Following extraction with a French press and centrifugation, soluble protein was subjected to ammonium sulfate precipitation and then purified on a Ni-NTA agarose column by washing in presence of 10 mM imidazole and elution with 250 mM imidazole. Protein was then further purified by Sephacryl S 300 size exclusion chromatography (GE Healthcare) and concentrated by ultrafiltration (Amicon Ultra-15, Merck-Millipore). The final buffer solution was 50 mM Tris/Cl, 5 mM EDTA, 300 mM NaCl, 10% glycerol, pH 7.8. Centricon Protein concentrations were measured according to Bradford [36].

#### Generation of (6-4) photoproducts

The oligonucleotide sequences are from [27], synthesized by Sigma-Aldrich, and in listed in Table S1. To obtain the (6-4) photoproduct, the oligonucleotide was dissolved in Millipore water at a concentration of 12.5 μM and degassed with argon. The solution was poured into a Petri dish; the thickness of the solution was 2 to 3 mm. The Petri dish was placed on a 4°C cooling pack in an irradiation box under argon atmosphere and irradiated with UV-C (GE Healthcare, G15T87B, 15 W) for 6 h at a distance of 12 cm. The irradiated DNA was concentrated by vacuum centrifugation and purified by HPLC on a “series 1200 Agilent Technologies system” using a Gemini C18 column (50 × 4.60 mm, 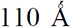, Phenomenex). Mobile phases were solution A (0.1 M TEAA in H_2_O) and solution B (0.1 M TEAA in H_2_O/ACN 20/80), the gradient was 4-18% B in 45 min, and the flow rate was set to 5 ml / min. The elution was monitored by recording UV/vis spectra every second. The (6-4) photoproduct is identified by an additional absorbance peak at 325 nm [28]. Relevant fractions of (6-4) photoproducts were collected and stored at −20 °C. The purity of (6-4) photoproducts was checked by subsequent analytic HPLC as described below for the repair assay. If the (6-4) photoproduct was not pure, a second HPLC purification followed under the conditions as described for the repair assay.

In order to obtain double stranded ODN5 oligonucleotides, (6-4) ODN5 and its complementary oligonucleotide (ODN5-Compl) were mixed at equimolar concentrations, heated to 95 °C for 3 min, and cooled to 25 °C over a period of ca. 90 min.

#### Repair assay

The 30 μL photorepair reaction mixture contained 5 μM of the (6-4) photoproducts in repair buffer (50 mM Tris-HCl, pH 7.5, 1 mM EDTA, 100 mM NaCl, 5 % (w/v) glycerol, 14 mM DTT). The protein concentration was 0.5 μM unless stated otherwise. In the repair assay with Mg^2+^, the concentration of free Mg^2+^ (after subtraction of the EDTA bound fraction) was 4 mM unless stated otherwise. After 5 min pre incubation in darkness at 22 °C, aliquots were irradiated with 400 nm light emitting diodes (250 μmol m^-2^ s^-1^) for the indicated time. After irradiation, EDTA was added into the reaction mixture to a final concentration of 8 mM to chelate the Mg^2+^. The reactions were stopped by heating to 95 °C for 10 min and centrifuged at 13,000 rpm for 10 min; the supernatants were subjected to HPLC. To this end, the above Agilent system was used with a Gemini C18 column (50 × 4.60 mm, 110 Å, Phenomenex). The HPLC conditions were [12]: 7 % ACN in 0.1 M TEAA (pH 7.0) for 0-5 min; 7-10 % ACN in 0.1 M TEAA (pH 7.0) for 5-35 min; flow rate, 0.75 ml/min; column temperature, 25°C for t_repair and 60°C in all other cases. The repair efficiency was estimated from the peak areas corresponding to damaged DNA and repaired DNA as described earlier [14].

#### Photolyase from *Prochlorococcus marinus*

PhrB homologs [6] were screened for sequences without FeS-binding cysteines. These formed a single clade in phylogenetic analysis. We selected a sequence from *Prochlorococcus marinus* ssp CCMP1986 [29] (Uniprot identifier or the protein: Q7V2P7) and synthesized a codon optimized gene (Genscript) for expression in the pET28a vector (Novagen). The expressed gene has an N-terminal His-tag. Protein was expressed in *E. coli* ER2566 in autoinduction medium [30] with 100 μM IPTG. The culture was cultivated for 3-4 d at 14 °C under 180 rpm shaking. Cell density reached OD_600 nm_ = 17-18. The extraction buffer was 40 mM Tris/Cl, 240 mM NaCl, 20% glycerol, 1 % Nonidet P-40, pH 7.4. Extraction and purification followed the PhrB protocol, but ammonium sulfate precipitation was omitted and for the binding to the nickel column the Ni-NTA resin (Qiagen) was shaken for 1 hour in the protein solution. The imidazol concentration of the washing and elution buffer was 40 mM and 250 mM, respectively.

#### Computational Methods

A PhrB-DNA complex model has been built using the PhrB WT crystallographic structure (PDB 4DJA) [6] and the damaged DNA bound to Drosophila melanogaster (6-4) photolyase (PDB 3CVU) [10]. Few residues have been slightly rotated to avoid severe steric clashes. Three starting structures have been defined considering the three possible protonation states of H366: Nε protonation, Nδ protonation or positively charged and double protonated histidine. The protonation state of other acidic and basic residues have been determined from pKa calculations using the PropKa webserver [37]. The force field parameters used for DMRL and the iron sulphur cluster have been described earlier [6]. The FADH^-^ parameters were taken from the AMBER force field [31, 32] and the charges were determined using the RESP approach [33].

All mentioned simulations were performed with the GROMACS 2016 package [34, 35] using the AMBER-SB99-ILDN force field [31, 32]. The protein-DNA complex was then solvated in a 103 Å side cubic water box and a combination of 21 Mg^2+^ and one Cl^-^ or 20 Mg^2+^ was randomly added in the box to ensure the global neutrality of the system in the presence of neutral or positively charged H366, respectively. An equilibration procedure consisting in a minimization, a 1 ns NVT simulation and a 1 ns NPT simulation with constrained bonds was performed.

Since the first model conformation of the protein-DNA complex is not stable in long MD simulations, constrained MD simulations were performed to obtain a better conformation of the damaged DNA in the active state and to allow equilibration of the interaction between the DNA and the protein surface. The 5’ O5-H366Nε and the 3’ N3-H366Nδ bonds have been both restrained to 3 Å using the pull plugin in GROMACS and a force constant of 1000 kJ·mol^-1^. The force of the constraint has been chosen as the criterion to define the convergence of the constrained MD simulations. 100 ns free classical MD simulations have been performed starting from the final structure of the corresponding constrained simulations and the stability of the complex has been tested by measuring distances between the DNA photoproduct and H366. Considering our results, the neutral H366 protonated on Nε appears less efficient in maintaining a stable complex between protein and DNA. Consequently, in the following only the neutral H366 protonated on Nδ, designated as neutral H366, and the positively charged H366^+^ have been considered. One microsecond NPT simulation has been performed considering these two protonation states of His-366.

The D254N and the D179N mutants were constructed from the final structure of the complex that resulted from the H366^+^ 100 ns free simulation. The DNA-protein complexes were solvated in a new water box and the same number of ions as for PhrB-WT were randomly added. Only the positively charged H366^+^ was considered in the mutants. The simulation boxes were equilibrated following the same procedure as for PhrB-WT. One microsecond NPT simulation has been performed for each mutant.

To determine the degree of conservation of the Lys246 and the Arg421 residues in eukaryotic (6-4) photolyases, we performed an alignment with the 1000 nearest BLAST (Uniprot Dec 2017) homologs of *Drosophila* 6-4 photolyase. At the position of Lys246 (of Drome 6-4 PL) there was a Lys in 42% and an Arg in 53% of all sequences. The remaining 5% sequences were annotated as animal cryptochromes or proteins with unknown functions. At the position of Arg421, 35% of the sequences also had an Arg residue, 57% had a His and 1% a Lys residue.

### Supplemental Tables

**Table S1.** Oligomers sequences. For repair assays, dissolved oligomers are irradiated with UV-C and the (6-4) photoproduct which is formed by the pair of thymines (bold) is purified by HPLC. Sequences are from [27]. The ODN5-Compl was used for double stranded DNA together with the photoproduct of ODN5.

| Name | Sequence (5'→3') | Length (nt) |
| --- | --- | --- |
| **t_repair** | AGG**TT**GGC | 8 |
| **ODN4** | CAGG**TT**GGCA | 10 |
| **ODN5** | ACAGG**TT**GGCAG | 12 |
| **ODN5-Compl** | CTGCCAACCTGT | 12 |
| **ODN6** | ACAGCGG**TT**GCAGGT | 15 |

**Table S2.** Repair of single stranded ODN5 by PhrB-Y424F. Mean values from 3 independent experiments ± SE. DNA and protein concentrations were 5 μM and 5 μM, respectively.

| | REA ($\text{min}^{-1}$) | fold increase |
| --- | --- | --- |
| no $\text{Mg}^{2+}$ | $(5 \pm 1) \times 10^{-4}$ | |
| with $\text{Mg}^{2+}$ | $0.2 \pm 0.005$ | 420 |

### Supplemental Figures

**Fig. S1.**
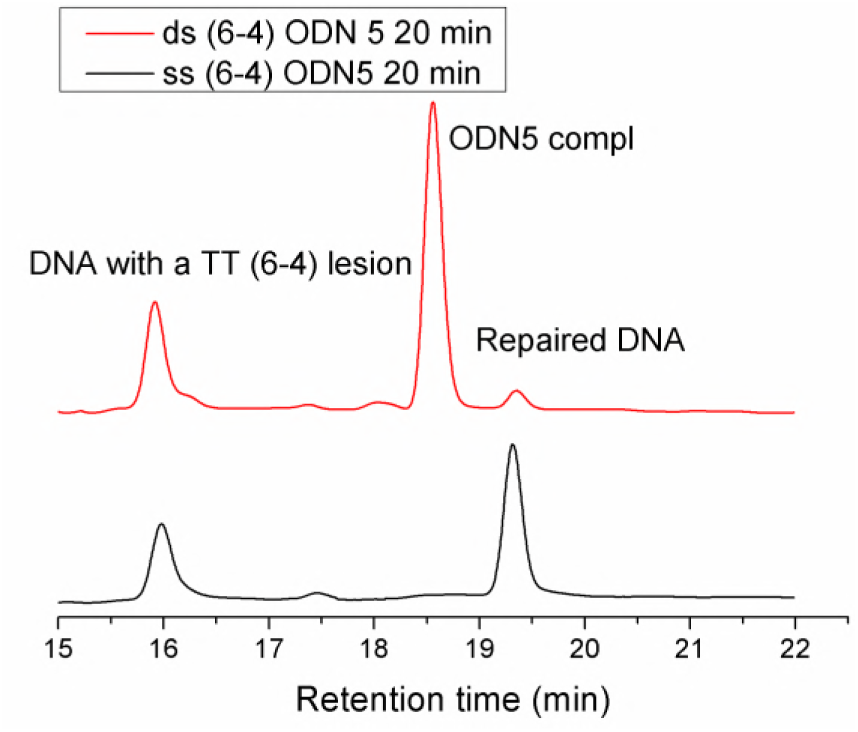
Examples of HPLC profiles of DNA repair by PhrB-WT: double stranded (6-4) ODN5 (12-mer) as substrate (upper red curve) and single stranded (6-4) ODN5 as substrate (lower, black curve), reaction mixture irradiated for 20 min in the absence of Mg^2+^.

**Fig. S2.**
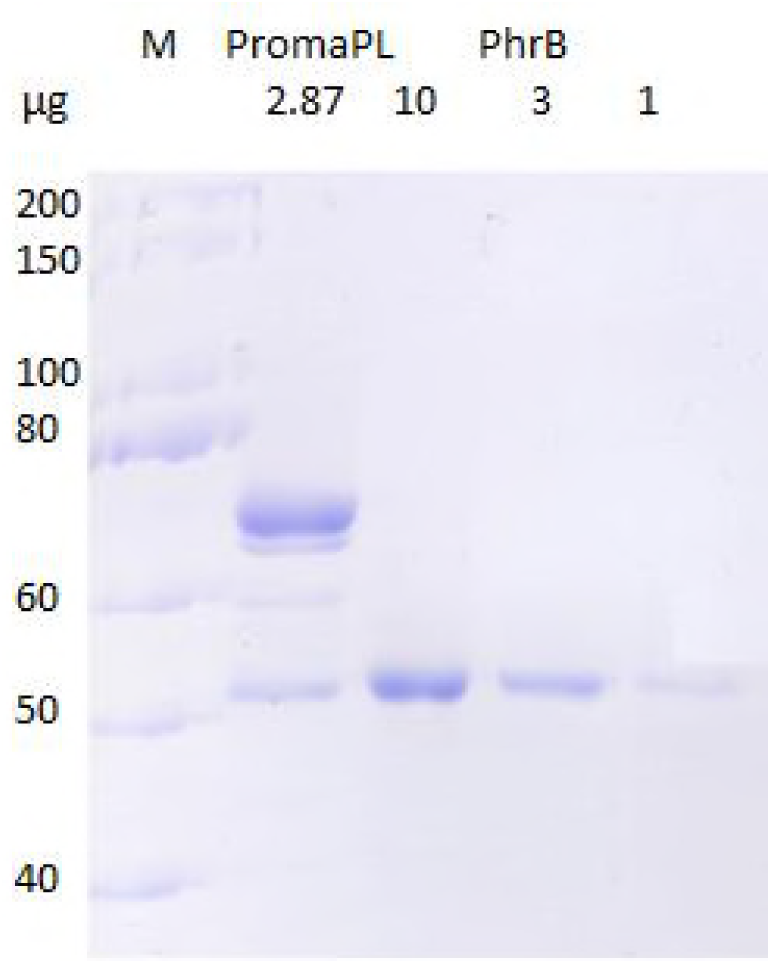
Coomassie stained SDS PAGE of purified PromaPL and varying amounts of PhrB. The size of protein markers in kDa is given on the left. Bands slightly above the 50 kDa marker refer to PromaPL and PhrB, respectively. The amounts of protein loaded are indicated above the gel, in the case of PromaPL the number was estimated from density values of a scanned image and refers to the specific band only.

